# Can suitability indices predict plant growth in the invaded range? The case of Acacias species

**DOI:** 10.1101/2022.12.15.520608

**Authors:** Carmen P. Silva, Daniela N. López, Paulette I. Naulin, Sergio A. Estay

## Abstract

Forestry in many parts of the world depends on exotic species, making this industry a source of invasions in some countries. Among others, plantations of the genus Pinus, Eucalyptus, Acacia, Populus, and Pseudotsuga underpin the forestry industry and are a vital component of many countries economies. Among woody plants, the cosmopolitan genus Acacia includes some of the most commonly planted trees worldwide. In order to prevent, manage and control invasive plant species, one of the most used tools is species distribution models. The output of these models can also be used to obtain information about population characteristics, such as spatial abundance patterns or species performance. Although ecological theory suggests a direct link between fitness and suitability, this link is often absent. The reasons behind the lack of this relationship are multiple. Chile is one of the countries where Acacia species, in particular, *A. dealbata* and *A. melanoxylon*, have become invaders. Here, we used climatic and edaphic variables to predict the potentially suitable habitats for *A. dealbata* and *A. melanoxylon* in continental Chile and evaluate if the suitability indices obtained from these models are associated with the observed performance of the trees along the country. Our models show that variable importance showed significant similarities between the variables that characterize each species’ niche. However, despite the high accuracy of our models, we did not observe an association between suitability and tree growth. This disconnection between suitability and performance can result from multiple causes, from structural limitations, like the lack of biotic interactions in the models, to methodological issues, like the usefulness of the performance metric used. Whatever the scenario, our results suggest that plans to control invasive species should be cautious in assuming this relationship in their design and consider other indicators such as species establishment success.

## Introduction

The vast majority of exotic species introductions are human-mediated, especially in the case of plants (Saul et al., 2017), where activities such as horticulture, agriculture, and forestry are among the main introduction pathways (Hulme et al., 2008). One of the most used tools for understanding the establishment of exotic species are species distribution models (SDMs). These models are intended to establish the environmental tolerance limits or habitat suitability for a particular species through the correlation of its known geographical distribution, i.e., occurrence/absence or abundance records, and the values of several environmental variables at the occurrence sites (Soberón and Peterson, 2005; Phillips et al., 2006; Elith and Leathwick, 2009; Lobo et al., 2010). The resulting environmental suitability estimates can also be used to obtain information about other population characteristics, such as spatial abundance patterns or species performance (Thornton and Peers, 2019). A high correlation between suitability and performance is desirable for several reasons. From a productive point of view, a strong relationship would facilitate the identification of the best sites for establishing planted forests for industrial purposes. Also, for invasive species control, a strong positive relationship would increase the probability of success of the control because the actions could be focused in locations where performance (or some proxy) is higher (Jarnevich et al., 2021). However, these models do not always show a strong relationship between suitability and performance (VanDerWal et al., 2009; Gutiérrez et al., 2013; Thuillier et al., 2010; Dallas and Hastings, 2018). Although ecological theory suggests a direct link between fitness (or some performance proxy) and suitability (Younginger et al., 2017), this link is absent in many cases. The reasons behind the lack of this relationship are multiple. On the one hand, if a species’ native distribution results from of dispersal limitations (e.g., insular species or with small global distributions), and not to the lack of tolerance to the environmental conditions in the new habitats, then suitability will be disconnected from performance. Some authors pointed out that this situation is equivalent to saying that the species’ phenotypic plasticity is greater than what may be appreciated from realized distributions (Orr and Smith, 1998; Qiao et al., 2017). On the other hand, the lack of association could be a consequence of the disconnection between the metric of performance used in the study and fitness. In particular, tree species planted worldwide for industrial purposes could show performances significantly different from those predicted by SDMs fitted using their native distribution.

Forestry in many parts of the world depends on exotic species, making this industry a source of invasions in some countries (Richardson, 1998). Among others, plantations of the genus *Pinus, Eucalyptus, Acacia, Populus*, and *Pseudotsuga* underpin the forestry industry and are a vital component of many countries’ economies (Richardson, 1998; Richardson and Rejmánek, 2011). Because of their economic value, information about their performance under several (and in many cases novel) climatic and edaphic conditions is available in many countries. This situation provides a unique opportunity to evaluate the relationship between suitability, obtained using information from the native distribution, and performance acquired using information from their non-native distribution.

Among woody plants, the cosmopolitan genus Acacia (*sensu lato*) (*Fabaceae*) includes some of the most commonly planted trees worldwide (Jansen and Kumschick, 2022), along with *Pinus* and *Eucalyptus* (Richardson et al., 2011). The genus Acacia *s.l* includes over 1,300 trees and shrubs found in Africa, Madagascar, Asia, and North and South America (Lorenzo et al., 2010), but most of them, 1,012 species, approximately, are native to Australia, collectively known as Australian acacias or wattles (Lorenzo et al., 2010; Miller et al., 2011). According to Richardson et al. (2011), as many as 386 Australian acacias have been introduced to areas outside their native ranges (Richardson et al., 2011), mainly because of their economic value and for restoration and ornamental purposes (Griffin et al., 2011; Kull et al., 2011). Currently, several Australian acacias are confirmed as invasive (Richardson et al., 2011; Wilson et al., 2011). One of them, *A. mearnsii*, is included in the “100 of the World’s Worst Invasive alien species” (Lowe et al., 2000), and *A. dealbata* is listed in the “100 of the worst invasive species in Europe” (Nentwig et al., 2017).

Chile is one of the countries where Acacia species, in particular, *A. dealbata* and *A. melanoxylon*, have become invaders (Langdon et al., 2019; Fuentes-Ramírez et al., 2011; Fuentes et al., 2014). Both species were initially introduced for ornamental and furniture manufacturing purposes, *A. dealbata* in 1869 and *A. melanoxylon* in 1923 (Fuentes-Ramírez et al., 2011; Fuentes et al., 2014), and their ranges seem to be still increasing (Langdon et al., 2019). Several studies address *A. dealbata* invasion in Chile and its impact on native vegetation(Fuentes-Ramírez et al., 2010; Fuentes-Ramírez et al., 2011; Pauchard and Maheu-Giroux 2007; Peña et al., 2007; Langdon et al., 2019). For *A. dealbata*, several SDMs have been developed to estimate its invasive potential, but only using climatic variables (Langdon et al., 2019; Bustamante et al., 2022). However, the association between the suitability obtained from these models and the actual performance of trees in the field has not been evaluated, so the design of control strategies using only these results could be based on highly uncertain scenarios. In the case of *A. melanoxylon*, no evaluation of its potential distribution has been performed. In this study, we estimate the potential distribution of *A. dealbata* and *A. melanoxylon* in Chile using climatic and edaphic variables and evaluate if the suitability indices obtained from these models are associated with the observed performance of the trees along the country.

## Methods

### Species occurrence data

Presence records of the native distribution of *Acacia dealbata* and *A. melanoxylon* were obtained from the Atlas of Living Australia (http://www.ala.org.au/) and the Global Biodiversity Information Facility (GBIF). Because our objective is to evaluate the relationship between suitability and performance, we only include data from the native distribution of both species. The original data sets were checked and filtered; all duplicate geographical records and those presenting incomplete or dubious information were deleted. To reduce geographical sampling bias, only one record in an area of ~1 km^2^ was considered. This process resulted in 11,683 points for *A. dealbata* and 18,146 for *A. melanoxylon*. We generate pseudo-absences using a 1:1 ratio following general recommendations (Valavi et al., 2021).

### Environmental layers

The environmental variables selected to implement SDMs are crucial; they directly impact the predictive accuracy and model realism (Mod et al., 2006). The variables should vary depending on the research question or the modeling goal (Irving et al., 2020; Araujo et al., 2019). For terrestrial plants, soil properties characteristics are essential, significantly impacting their establishment and growth, thus influencing their distribution (Beauregard and de Bios 2014). Hence, incorporating edaphic factors is desirable and may improve model performance and enhance the accuracy of the outcome (Coudun et al., 2006).

Here, we used climatic and edaphic variables to predict the potentially suitable habitats for *A. dealbata* and *A. melanoxylon* in continental Chile. Current climatic conditions were obtained from the Chelsa database (Karger et al., 2017), while soil variables were gathered from the Global Soil Dataset (Shangguan et al., 2014), with a spatial resolution of 30 arcsec. Edaphic layers are available at depths from 0 to 2.3 m, but layers between 0 and 1.4 m are highly correlated (*r* >0.9). For this reason, our analysis was performed using the layers corresponding to depths between 5 to 19 cm. Initially, a preselecting variables approach to avoid the risk of multicollinearity, based on the species biology, was applied; climatic and edaphic predictors were analyzed together. We eliminated the predictor variables yielding correlation values above 0.7 (Pearson’s coefficient) in the pairwise cross-correlation matrix or those with apparent unclear biological importance. The final sets of used variables are shown in Table 1. The training area was defined using a buffer of 500 km around presence points. The buffer size was determined considering an approximation to the geographic area accessible to the species in a time covering several generations (Barve et al., 2011).

**Table 1:** Variables used in the study

| Code | Description | Units |
| --- | --- | --- |
| BIO1 | mean annual air temperature | °C |
| BIO10 | mean daily mean air temperatures of the warmest quarter | °C |
| BIO11 | mean daily mean air temperatures of the coldest quarter | °C |
| BIO12 | annual precipitation amount | Kg m<sup>-2</sup> year<sup>-1</sup> |
| ALT | altitude | masl |
| TC | total carbon | % of weight |
| TN | total nitrogen | % of weight |
| TP | total phosphorus | % of weight |

### Modeling approach

We used the Regularized Random Forest algorithm (RRF, Deng and Runger, 2012) to estimate niche models for *A. dealbata* and *A. melanoxylon* using the R package RRF. This algorithm uses a regularization process to discard the least important variables, producing more parsimonious models, with a similar prediction error to the full model (Deng and Runger, 2012). A regularization coefficient was applied using the scheme proposed by Deng (2013) for Guided Random Forest with a γ = 1 for the maximum penalty. The hyper-parameter mtry was defined using the function tuneRRF in the package RRF. We used a 5-fold cross-validation scheme for each model and divided our dataset in 70/30 for training and testing subsets. We set the Ntree hyper-parameter in 1000. We evaluated them using the area under the curve (AUC) of the ROC curve, True Skill Statistics (TSS), and Symmetric Extremal Dependence Index (SEDI) using the test data subset. Variable importance was evaluated using the mean decrease Gini. As a complement to evaluate the similarities of species niche models, we calculate the overlap between the hypervolume described by the ellipsoid that represents the environmental niche of both species using the package SIBER (Jackson et al., 2011). All analyses were performed in R 4.2.1 (R Core Team 2022).

### Suitability – Performance relationship

The final step was to evaluate if the modeled suitability of *A. dealbata* and *A. melanoxylon* in Chile is correlated to the species’ performance. We gathered the data from the report “Progress in research with species of the genus Acacia in Chile” (Pinilla et al., 2010). The main goal of this study was to assess which species show the highest growth under different edaphic and climatic conditions in Chile.

To estimate *A. melanoxylon* and *A. dealbata* performance in Chilean territory, we calibrate a logarithmic height-age (height ~ Ln(age)) curve using all data available from the field trials (38 sites for *A. dealbata* and 31 sites for *A. melanoxylon)*, obtaining a function that estimates the height of the trees as a function of age. We took the function’s standardized residual as a proxy of how much or less the trees in the site grow over the expected value (observed performance). At each site, we took the suitability median values in a buffer of 2,500 meters around the coordinates of the trial. To evaluate the relationship between suitability values and the observed performance, we follow the recommendation of VanDerWal et al. (2009). They suggest that suitability indices are more associated with the maximum potential performance than with the average performance. In this vein, we use linear regression and linear quantile regression (90% percentile) to determine if suitability indexes successfully predict the observed performance of both species. We assessed the magnitude of the relationship using the 95% confidence interval of the slope of each regression.

## Results

### Models Performance

Regularized Random Forest models showed high predictive accuracy for *A. dealbata* and *A. melanoxylon*. Performance measures are given in Table 2. Model projections show that *A. dealbata* showed moderate suitability in southern Chile, while *A. melanoxylon* indicates moderate to high suitability in central-south Chile (Fig. 1). An analysis of variable importance showed differences between species (Table 3). Mean annual air temperature (BIO1), annual precipitation amount (BIO12), and altitude (ALT) were the top three important variables for *A. dealbata*. While mean daily mean air temperature of the warmest quarter (BIO10), annual precipitation amount (BIO12), and altitude (ALT) were the most important variables for *A. melanoxylon*. Considering the overlap between ellipsoids, 97% of the environmental niche of *A. dealbata* is contained inside the environmental niche of *A. melanoxylon*. However, only 67% of the niche of the latter is contained in the niche of the former (Fig. 2).

**Table 2:** Accuracy metrics for SDMs. See methods for details.

| Species | Accuracy | Sensitivity | Specificity | AUC | TSS | SEDI |
| --- | --- | --- | --- | --- | --- | --- |
| *A. melanoxylon* | 0.915 | 0.941 | 0.892 | 0.968 | 0.833 | 0.933 |
| *A. dealbata* | 0.904 | 0.937 | 0.876 | 0.957 | 0.813 | 0.922 |

**Figure 1:**
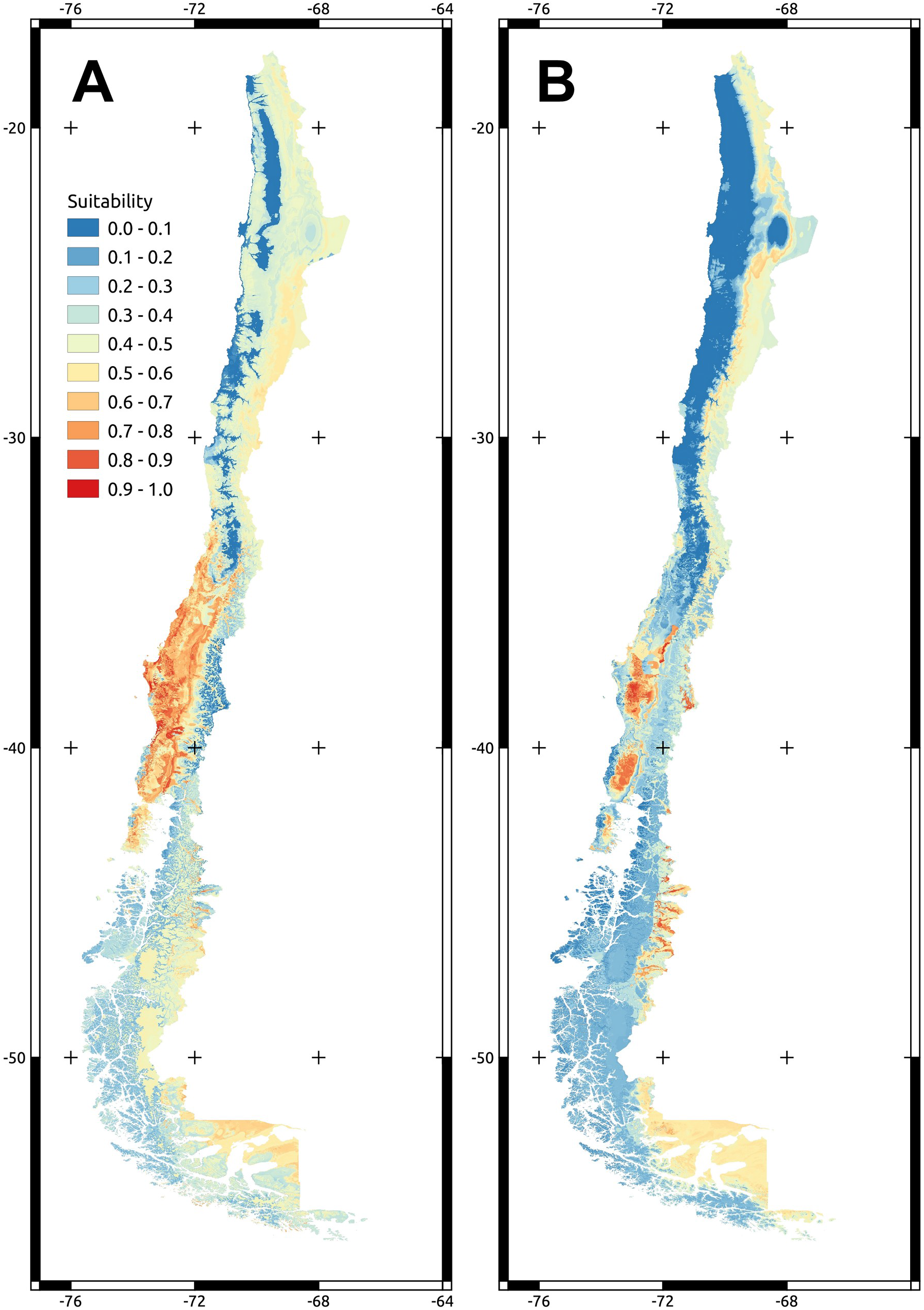
Maps showing projected models of A) *A. melanoxylon* and B) *A. dealbata* over continental Chilean territory.

**Table 3:** Relative importance of variables on SDMs for *A melanoxylon* and *A. dealbata*. Importance is expressed as the mean decrease Gini index.

| *A. melanoxylon* | Importance | *A. dealbata* | Importance |
| --- | --- | --- | --- |
| BIO10 | 100.00 | BIO1 | 100.00 |
| **BIO12** | **15.02** | **BIO12** | **20.81** |
| ALT | 10.30 | ALT | 19.08 |
| **BIO1** | **6.63** | **BIO11** | **15.43** |
| BIO11 | 5.24 | BIO10 | 9.42 |
| **TC** | **1.76** | **TC** | **3.34** |
| TN | 0.66 | TN | 2.85 |
| **TP** | **0.00** | **TP** | **0.00** |

**Figure 2:**
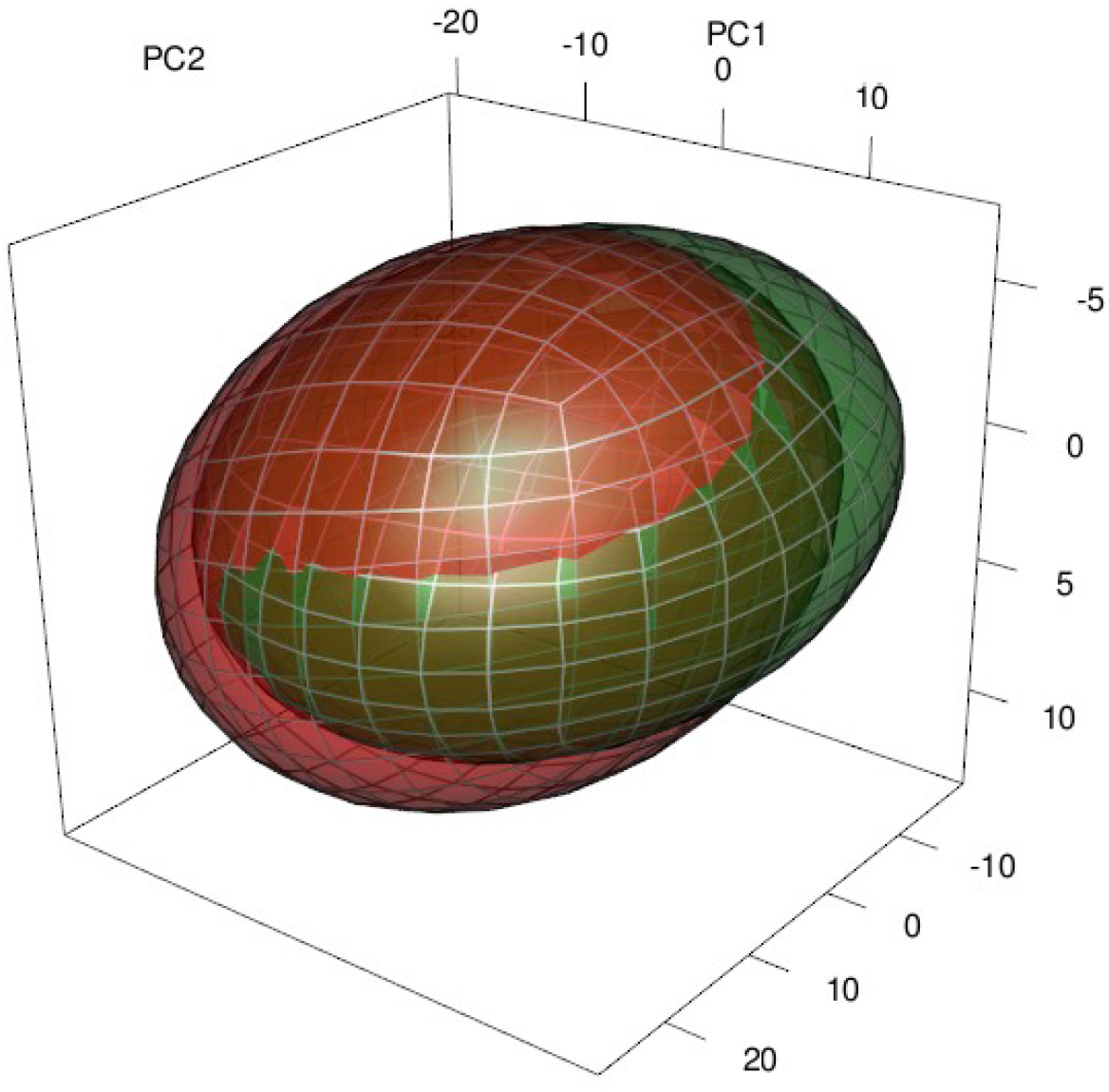
Representation of the environmental niches of both species through ellipsoids and the overlap between them. The 3-D space corresponds to the three first principal components calculated with the same environmental variables used in the niche models. In red *A. melanoxylon*, green *A. dealbata*.

None of the edaphic variables included in the analysis were identified as important in the final models.

### Suitability - Performance relationship

The goodness of fit of all models was very low (R^2^ and McFadden’s pseudo R^2^ values, see Table 4). The relationship between suitability and performance was weak for all regressions. For both species, the value of the slopes was non-significant according to the 95% confidence intervals (Table 4).

**Table 4:** Results for simple lineal and quantile lineal models for the relationship between sustainability and tree growth for both Acacia species.

| Linear model | Intercept | Slope | Slope 95% CI | R2 |
| --- | --- | --- | --- | --- |
| *A. melanoxylon* | 0.749 | -1.035 | [-3.744, 5.243] | 0.004 |
| *A. dealbata* | -0.44 | 0.624 | [-1.644, 0.765] | 0.013 |

| Quantile lineal model | Intercept | Slope | Slope 95% CI | Pseudo-R2 |
| --- | --- | --- | --- | --- |
| *A. melanoxylon* | -1.757 | 3.969 | [-12.46, 22.43] | 0.039 |
| *A. dealbata* | 1.119 | 0.107 | [-3.493, 2.774] | 0.003 |

## Discussion

We obtained a good model performance for *A. dealbata* and *A. melanoxylon*. Although both species presented areas with moderate to high suitability south of 33° S (Fig. 1), the latter has a larger area of high suitability. The evaluation of variable importance showed significant similarities between the variables that characterize each species’ niche. These results were expected considering the native habitats of these species. The native distribution range of *A. dealbata* is narrower than *A. melanoxylon. Acacia melanoxylon* occupies well-drained soils in cool and warm, humid climates (Weber 2017; CABI 2022). Maximum and minimum temperatures in the native range from 23 to 26° C and 1- 10°, respectively (Weber, 2017; CABI, 2022). *Acacia dealbata* grows under drier conditions on several soil classes in cool to warm sub-humid climates (Weber, 2017; CABI, 2022). This species occupies habitats with over 500 mm rainfall, usually at altitudes from 350–1000 m above sea level (May and Attiwill, 2003; Lorenzo et al., 2010). Both species are fast-growth colonizers that can expand their initial introduction range by establishing new populations, usually associated with rivers, roads, post-fire, and degraded lands, i.e., strongly associated with anthropogenic disturbances (Matthei, 1995; Pauchard and Maheu-Giroux, 2007; Peña et al., 2007). Both species are problematic in Chile, and caution is advised in their silvicultural management (Peña et al., 2007; Langdon et al., 2019). According to our results, the main concern is the potential expansion of both species southward of their current limit, similar to previous results (Langdon et al., 2019). Fuentes et al. (2014) indicated that the current southern limit of both species in Chile occurs near 43°S. Our projections show that the most suitable habitats for both species are far south of this limit (Southern Patagonia), which adds our results to the several calls to increase control efforts to prevent colonization beyond the current invaded area (Langdon et al., 2019).

The fact that none of the edaphic variables were important when characterizing the species’ environmental requirements may be explained by the fact that both *A. dealbata* and *A. melanoxylon* can fix atmospheric nitrogen (Quiroz et al., 2009; Brockwell et al., 2000). In South Africa, Gouws & Shackleton (2019) also found no relationship between soil properties and plant density or biomass. *Acacia dealbata* increases several nutrient concentrations (Lorenzo et al., 2010). Potentially available nitrogen, total nitrogen, and organic carbon increase in habitats with *A. dealbata* (May and Attiwill, 2003; Lorenzo et al., 2010), with long terms effects (Souza-Alonso et al., 2015).

Despite the high accuracy of our models, we did not observe an association between suitability and tree growth. Midolo et al. (2021) suggested that an implicit and scarcely tested assumption in niche models is that individual fitness should be higher at the center of the environmental niche, what they called the “fitness-centre” hypothesis. However, they found that the support for this hypothesis in actual data was scarce (Midolo et al., 2021). Similarly, Bernal-Escobar et al. (2022) said that, according to the fitness-suitability hypothesis, there should be a positive relationship between climate suitability and tree growth rates. These authors, however, found a negative relationship in both gymnosperms and angiosperms trees. In a more detailed analysis, Sanchez-Martinez et al. (2021) suggested that the positive relationship between suitability and tree growth exists, but only for models fitted using locations with the highest performance (top 10-30% tree growth). In this sense, the relationship seems valid only for sites where trees show very high performance (Sanchez-Martínez et al., 2021). However, including sites where growth is moderate or low weakens the relationship. A potential explanation comes from the inclusion in the dataset of sink populations outside the fundamental niche, where fitness is null, and the species occurrence depends exclusively on propagule arrival (Guisan et al., 2017). Also, in the particular cases of *A. dealbata* and *A. melanoxylon*, these species presents high plasticity to soil water availability and other environmental conditions (Pohlman et al., 2005) and is capable of modifying from soil chemical properties to soil and plant microbial communities (Lorenzo et al., 2010; González-Muñoz et al., 2012; Lazzaro et al., 2014; Guisande-Collazo et al., 2016), a situation that has also been confirmed in Chile (García et al., 2012). This capacity is a critical factor that makes this species a successful invader since these soil modifications boost the establishment of its seedlings (Lorenzo et al., 2017).

Dolos et al. (2015) proposed a different explanation. Since suitability indexes do not consider the influence of pathogens/herbivores and competition on species distribution and their influence on mortality, including demographic information and interactions (mortality, herbivory, among others) may significantly improve these models’ performance (Dolos et al., 2015). For example, Acacia invasions usually take advantage of human-mediated disturbances (Lorenzo et al., 2010). In central-south Chile, Acacias could be excluded from some suitable sites due to plant community resistance. However the occurrence of removal of native flora or wildfires provides the opportunity for colonization. In these sites, the relationship between suitability and performance is absent under the lack of disturbances, but after these sudden changes, the relationship emerges.

Despite all these explanations, the lack of association may occur because plant growth is not a good proxy for fitness or because fitness depends on biophysical factors different from those used in training the SDM (Bernal-Escobar et al., 2022).

The relationship between suitability and performance has been reviewed mainly using abundance in other groups (e.g., VanDerWal et al., 2009; Januario et al., 2015). In plants, Weber et al. (2017) reported correlations between suitability and abundance mostly below 0.4 (Fig 3A in Weber et al., 2017). On the other hand, Dallas and Hastings (2018) report that suitability is mostly unconnected to abundance after trained models for 158 species, showing correlations close to zero in most cases. Patch size and plant dispersal limitations have been suggested as potential factors causing this lack of association (Dallas and Hastings, 2018). However, these factors are irrelevant for forest plantations, where site, plantation density, and management are human-mediated.

For many reasons, a strong correlation between suitability and performance is desirable and theoretically plausible. Control, like invasive species management (Jarnevich et al., 2021) or conservation activities (Midolo et al., 2021), would benefit from SDMs with a strong association with performance; however, our results and several others pointed out a weak association in most real cases. In this context, plans to control invasive species should be cautious in assuming this relationship in their design and consider other indicators such as species establishment success.

## Acknowledgments

The authors were supported by ANID PIA/BASAL FB0002 and Fondecyt 1211114

